# Periodicity Scoring of Time Series Encodes Dynamical Behavior of the Tumor Suppressor p53

**DOI:** 10.1101/2020.02.04.933192

**Authors:** Caroline Moosmüller, Christopher J. Tralie, Mahdi Kooshkbaghi, Zehor Belkhatir, Maryam Pouryahya, José Reyes, Joseph O. Deasy, Allen R. Tannenbaum, Ioannis G. Kevrekidis

## Abstract

In this paper, we analyze the dynamical behavior of the tumor suppressor protein p53, an essential player in the cellular stress response, which prevents a cell from dividing if severe DNA damage is present. When this response system is malfunctioning, e.g. due to mutations in p53, uncontrolled cell proliferation may lead to the development of cancer. Understanding the behavior of p53 is thus crucial to prevent its failing. It has been shown in various experiments that periodicity of the p53 signal is one of the main descriptors of its dynamics, and that its pulsing behavior (regular vs. spontaneous) indicates the level and type of cellular stress. In the present work, we introduce an algorithm to score the local periodicity of a given time series (such as the p53 signal), which we call *Detrended Autocorrelation Periodicity Scoring* (DAPS). It applies pitch detection (via autocorrelation) on sliding windows of the entire time series to describe the overall periodicity by a distribution of localized pitch scores. We apply DAPS to the p53 time series obtained from single cell experiments and establish a correlation between the periodicity scoring of a cell’s p53 signal and the number of cell division events. In particular, we show that high periodicity scoring of p53 is correlated to a low number of cell divisions and vice versa. We show similar results with a more computationally intensive state-of-the-art periodicity scoring algorithm based on topology known as Sw1PerS. This correlation has two major implications: It demonstrates that periodicity scoring of the p53 signal is a good descriptor for cellular stress, and it connects the high variability of p53 periodicity observed in cell populations to the variability in the number of cell division events.

**AMS subject classification:** 92C42, 92C37, 62M10

## 1. INTRODUCTION

The tumor suppressor p53 is one of the most studied proteins due to its importance in preventing cancer, see e.g. Lahav (2009); Levine (1997, 2020); Vogelstein et al. (2000) for an overview. p53 is activated by cellular stress, such as DNA damage. As a consequence, the stress response pathway is initiated, leading to cellular outcomes such as DNA repair, cell-cycle arrest, or apoptosis (cell death). Malfunction of p53 can leave DNA damage undetected, causing uncontrolled cell proliferation, and subsequently, cancer.

It has been shown that in more than half of all human cancers the p53 gene is mutated, and that the associated protein is frequently inactive (Levine (1997); Jin and Levine (2001)). In order to detect and prevent p53 malfunction, it is thus important to understand its behavior under both stressed and unstressed conditions.

Many studies have been devoted to the functionality and behavior of p53, see Levine (2020) and references therein. In this paper we are interested in one important aspect, namely, p53’s long-term (5 day) dynamics in single cells.

It has been demonstrated in multiple experiments that p53 shows oscillatory behavior when exposed to stress such as gamma radiation (for example, see Geva-Zatorsky et al. (2006); Lahav et al. (2004); Reyes et al. (2018); Loewer et al. (2013)), but the dynamics might vary across tissues (Stewart-Ornstein and Lahav (2017)). The observed oscillations were reported to be either damped or periodic in cell populations, and they were found to be periodic only in single cells. The observed damped oscillatory behavior was therefore a result of averaging over many cells in a population (Lahav et al. (2004); Lahav (2009)). Furthermore, single-cell experiments (e.g., Lahav et al. (2004); Reyes et al. (2018)) show that the pulses of p53’s periodic oscillations become more regular when the cell is exposed to more radiation, which, as a result, increases the amount of DNA damage.

The induction of p53 under an exterior stress (such as radiation) is experimentally well-studied, and explained via a negative feedback loop with Mdm2 (Geva-Zatorsky et al. (2006); Lev Bar-Or et al. (2000); Ma et al. (2005)). Under basal (i.e., unstressed) conditions, however, p53 was believed to stay at a low level —an assumption again based on averaging over cell populations (Lahav (2009); Loewer et al. (2010); Michael and Oren (2003)). The single cell experiments of Loewer et al. (2010) show that proliferating cells exhibit spontaneous pulses in p53, which are connected to intrinsic cell damage during normal growth. They further explain that a cell can distinguish between modest damage (e.g., due to growth) and sustained damage (e.g., due to radiation). Cells with spontaneous pulsing behavior in p53 do not activate the stress response pathway. All of the discussed experiments and results show that periodicity of the p53 signal is one of the main descriptors of its dynamics, and that the pulsing behavior (e.g., regular or spontaneous) further indicates the level and type of cellular stress.

One of the main tools to analyze periodicity of time series is the *autocorrelation function* (ACF). It provides information about the internal structure of a time series signal by measuring the correlation (similarity), if any, between data points in the signal and their shifted counterparts, see Box and Jenkins (1976). The ACF can provide some interesting insights on time series, e.g., randomness, short-term correlation, non-stationarity, and periodic fluctuations, cf. Chatfield (2003). Different algorithms, based on the ACF, have been proposed to detect periodic variations in time series, e.g., Martin and Mailhes (2010) and Vlachos et al. (2005). Regularity of periodicity is often analyzed with pitch detection, through the autocorrelation function (see Box and Jenkins (1976)), and used to analyze p53 signals in e.g., Geva-Zatorsky et al. (2006). Via autocorrelation, a value between 0 and 1 (called the *pitch score*) is assigned to a given time series, where the value 1 means maximal periodicity. For example, the sine curve has a pitch score equal to 1. In the present work, we propose an algorithm, called *Detrended Autocorrelation Periodicity Scoring* (DAPS), which is localized version of the pitch detection algorithm, to analyze the dynamical behavior of single cell p53 time series.

The remainder of this paper is organized as follows. In Section 2, we describe the main components of the proposed algorithm for general time series analysis, and, in particular, for single-cell p53 time series. We start by briefly introducing sliding window embeddings in Section 2.1. Then, the ACF and the pitch score are defined in Section 2.2, before describing the main steps of the proposed localized version of the pitch detection algorithm DAPS in Section 2.3. The latter applies pitch detection to shorter sliding windows of a normalized version of the original time series, resulting in a *distribution of pitch scores*, rather than a single pitch score. To empirically validate DAPS, we use the state-of-the-art periodicity scoring algorithm Sw1Pers (Sliding Window 1-Persistence Scoring) of Perea and Harer (2015), which is briefly described in Section 2.4. We further introduce the AUROC (area under ROC curve) in Section 2.5, which is used to statistically test our hypotheses.

Next in Section 3.1, we argue that the pitch score distributions produced by DAPS give a more refined description of the overall periodic behavior of p53 signals. Indeed, localization is particularly advantageous in cases where the time series shows different periodicity behavior in the course of a single experiment. Change of periodic behavior is observed in long-term experiments (> 1 day), as the cell usually undergoes more than one cell cycle. In Section 3.2, we study the long-term (5 days) basal behavior of p53 in single cells using DAPS. While Loewer et al. (2010) describe the basal dynamics in single cells (traced over 1 − 2 days) by spontaneous pulsing (as opposed to regular pulsing), it is reported that, to their surprise, a fair amount of unstressed cells do actually show regular pulsing behavior similar to the one of irradiated cells. A similar behavior of p53 in cells under basal conditions is observed in the experiments of Reyes et al. (2018). We show a correlation between p53 periodicity (computed via DAPS) and the number of cell division events ^1^. In particular, we show that high periodicity scoring is correlated to low number of cell divisions, and vice versa. This implies that a cell population exhibiting variability in the number of cell divisions will also have variable p53 periodicity across the population.

The connection between number of cell divisions and p53 periodicity can also be rationalized from a biological point of view: High periodicity scoring translates to p53 activity, which stops cells from dividing, allowing time for DNA repair and preventing the expansion of damaged cells. Naturally, in the case of cell death, the cell stops dividing as well. However, the number of cell deaths encountered in the experimental data we analyze was negligible. Under basal conditions, there is no targeted exterior stress imposed on the cells (such as radiation), which would explain high p53 activity. It is nevertheless possible that cellular stress is present due to other (unknown) internal or external sources leading to p53 activation.

## 2. LOCALIZED PERIODICITY SCORING

We now present details of the two tools we use to score periodicity in time series. In the discussion below, we consider a sequence of time-series data points 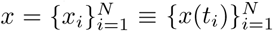 of length *N*. In our application, the time series is sampled once every 30 minutes, and *N* = 240, for a total of 5 days.

### 2.1 Sliding Window Embeddings

Before we introduce the tools, we first present a preprocessing step common to both. The *M-dimensional sliding window embedding* transforms a 1D time series into a sequence of *M*-dimensional Euclidean vectors as follows

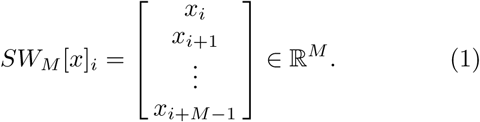

This transforms a time series into a point cloud. For instance, periodic time series turn into sampled loops, which we exploit in our analysis. More generally, the sliding window embedding stems from a theoretical result that allows one to reconstruct a state space of a dynamical system from a single observable, cf. Takens (1981), and it has been used in myriad applications, including analysis of vocal fold disorders (Herzel et al. (1994)), EEG analysis (Stam (2005)), activity recognition (Frank et al. (2010); Venkataraman and Turaga (2016)), and music information retrieval (Bello (2011)).

### 2.2 Autocorrelation function

Let the mean value of a time series *x* be denoted by 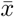. The ACF for lag *τ* is defined as follows:

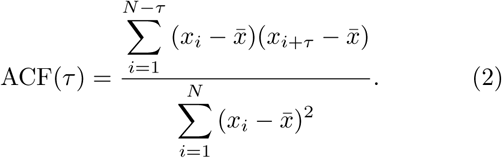

We also define the *pitch score* as the highest local maximum of the corresponding ACF

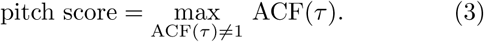

The pitch score is a value between 0 and 1, where 1 means maximal periodicity. The higher the pitch score, the more regular the pulses are. For example, a pitch score of 1 is assigned to a sine curve.

### 2.3 DAPS - Detrended Autocorrelation Periodicity Scoring

We now describe the algorithm we propose in this study, called Detrended Autocorrelation Periodicity Scoring (DAPS). The first step is a detrending and amplitude normalization using the sliding window embedding, which is performed as follows:

1. Given the sliding window of length *M* of an *N* -length time series *x*, arrange all of the sliding windows into the columns of an *M* × (*N* − *M* + 1) matrix *X*, so that the *i*^th^ column of *X* is *SW*_*M*_ [*x*]_*i*_. We note that *X* is a Hankel matrix; that is *X*_*i,j*_ = *X*_*i*+*k,j*−*k*_ for all integer *k* so that 1 ≤ *i* + *k* ≤ *M* and 1 ≤ *j* − *k* ≤ *N* − *M* + 1

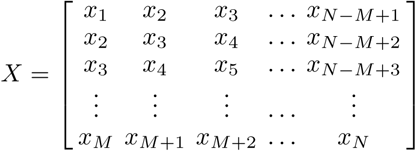 In other words, the *i*^*th*^ skew-diagonal of *X* is a constant value equal to the *i*^th^ value of the time series *xi*. We will exploit this observation in step 4. In the p53 application, we let *M* = 11, or 5.5 hours, which is roughly the length of one period of p53, cf. Geva-Zatorsky et al. (2006).
2. Subtract off the mean of each column of *X* from that column. This will normalize for linear drift in the time series, and it is referred to as “point centering” by Perea and Harer (2015).
3. After point-centering, normalize each column to have unit norm. This is meant to control for changes in amplitude, and is referred to as “sphere normalization” by Perea and Harer (2015). We now refer to the point-centered/sphere-normalized sliding window matrix as 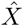.
4. At this stage, we deviate from Perea and Harer (2015). While they directly analyze the transformed point cloud represented by the matrix 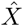 (Section 2.4), we instead solve for a time series whose sliding window is as close as possible to 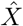, which should look like a version of *x* that is detrended. After the normalizations in steps 2 and 3, however, 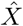 is no longer a Hankel matrix, which is a necessary condition for a sliding window matrix of a time series, as observed in step 1. Hence, we project 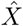 onto the space of Hankel matrices (under the Frobenius metric) by creating a new matrix *Y*, so that

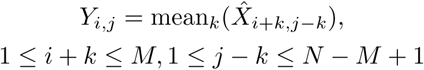 Expressed in simpler terms, *Y* is merely Hankel matrix in which each element is the mean of its corresponding skew-diagonal in 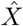. We can now read-off a time series from *Y* as the value of each successive skew diagonal. We refer to this final, detrended time series as *y*.

A simple example of the detrending is warranted at this point. Consider the first 50 samples of the time series

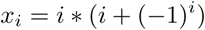

This is a linearly drifting and linearly amplitude modulated signal with oscillations hidden inside of it, and Figure 1 shows the detrending steps applied to this signal for a sliding window of size *M* = 5, which results in a periodic signal without drift (up to edge effects).

**Fig. 1.**
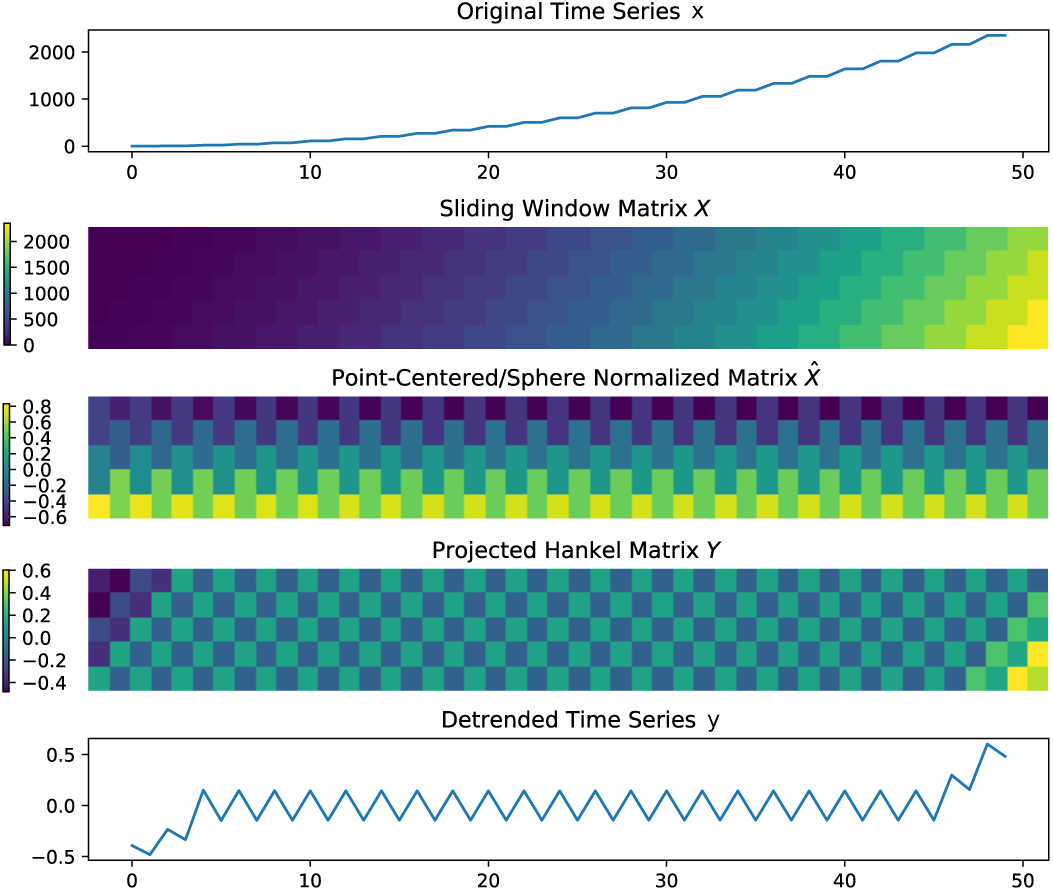
The detrending steps of DAPS allow us to extract periodic portions of a signal that is undergoing drift and amplitude variations.

Now that we have a detrended time series, we pull out every 24 hour contiguous chunk of *y* (48 samples), which we refer to as a “block” of *y*, and we compute a pitch score (Equation 3) for that block, which completes the DAPS algorithm. Based on the computed scores in each window, we obtain a distribution that reflects the local periodicity of the entire time series. A summary of the DAPS algorithm is outlined in Figure 2.

**Fig. 2.**
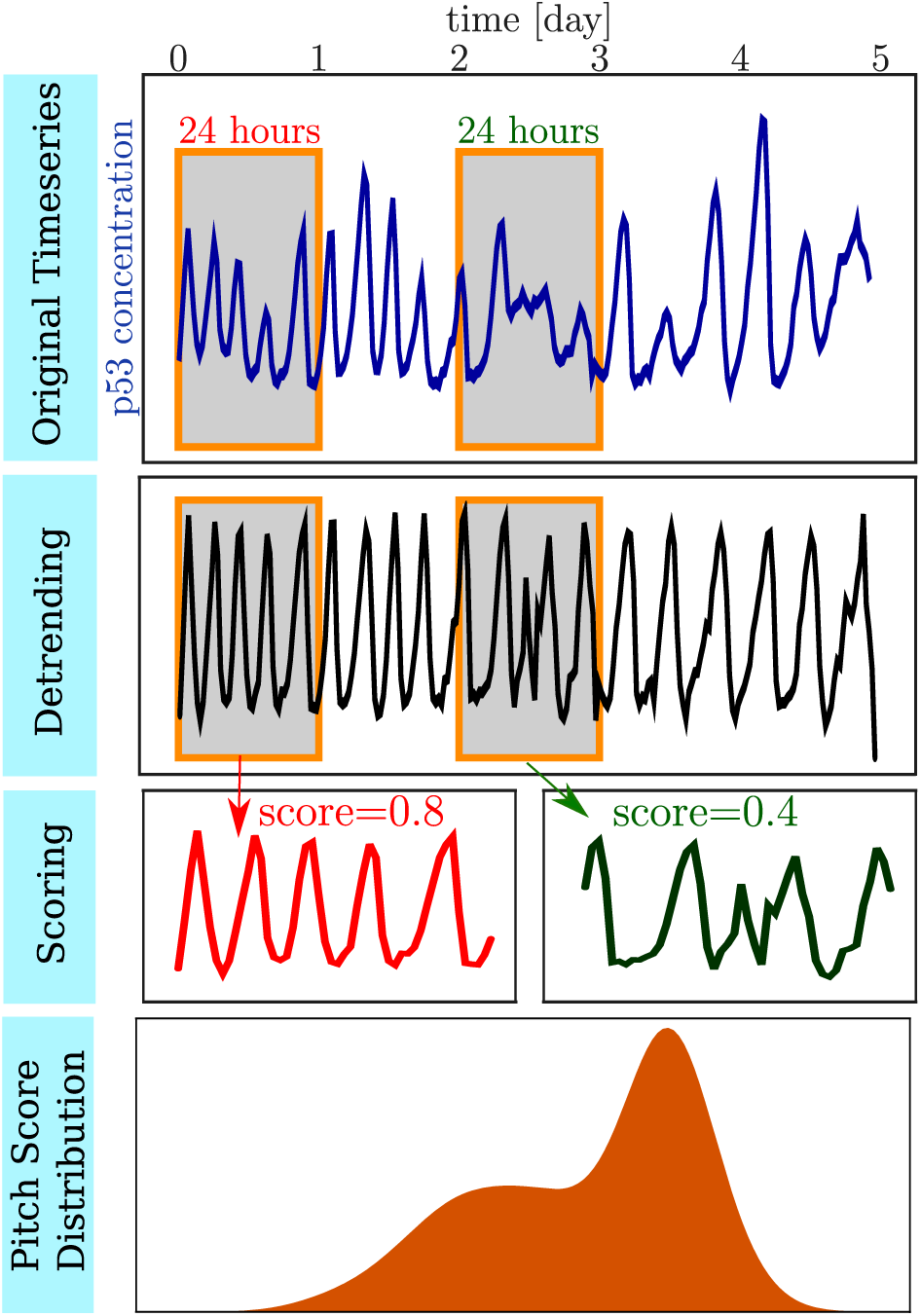
Schematic summary of the DAPS algorithm (Section 2.3) applied to a p53 time series. A time series is assigned a periodicity score distribution via the following steps: (i) Normalization (detrending) to have zero mean and unit norm; (ii) Splitting of time series into shorter intervals through a sliding window (in the case of p53, a window of length 24 hours is used, since this is the approximate length of a cell cycle); (iii) Computation of the pitch score in each 24 hour interval; and (iv) Summary of all pitch scores in a distribution.

Note that the DAPS algorithm assigns a distribution of periodicity scores to a time series, instead of a single value (which would be obtained by computing the pitch score of the entire times series only). The detrending with sliding windows is advantageous if the time series changes its periodicity behavior over time–a phenomenon encountered in biological times series, such as p53 (see the experimental results reported in e.g., Geva-Zatorsky et al. (2006); Loewer et al. (2013); Reyes et al. (2018)).

Furthermore, by summarizing the periodicity scores over different sliding windows in a pitch score distribution, we analyze the local behavior and sidestep global synchronization problems that result from an unknown internal biological time of each cell. In particular, we cannot compare two measurements of p53 directly to one another, since the time at the first sample of a time series *A* is not the same as the time at the first sample of another time series *B*, because cell *A* might just have divided, but cell *B* is shortly before a division. To control for this globally, one would have to synchronize the cells with respect to the cell cycle (which is done in some experiments in e.g., Reyes et al. (2018) or Loewer et al. (2010)), but this is a moot point with our procedure.

### 2.4 Comparison to Sw1PerS

To empirically validate DAPS, we compare it to a state-of-the-art algorithm for quantifying periodicity known as *Sliding Window 1-Persistence Scoring* (Sw1PerS) (Perea and Harer (2015)), variants of which have been used to detect genes regulating circadian rhythms (Perea et al. (2015)), turning and chatter in mechanical systems (Khasawneh and Munch (2014, 2016)), and wheezing in audio Emrani et al. (2014), for instance. As with DAPS, Sw1PerS starts off with a sliding window embedding, followed by point-centering and sphere normalization for detrending, but it uses techniques from topological data analysis (Edelsbrunner and Harer (2010)) to measure the “circularity” of resulting windows in ℝ^*M*^. A perfect sine wave with an *M* matching its period will give rise to a perfect circle in the sliding window embedding, which will receive a score of 1. Signals with imperfect repetitions will receive lower scores, with the lowest possible score being 0. As with DAPS, we use a sliding window of length *M* = 11. For technical reasons related to “birth time” of the sliding window loop (see Perea and Harer (2015)), we also upsample the sliding window point cloud so that there are 5× as many windows (see spline interpolation steps in Perea and Harer (2015) for more details). As we show in Section 3, Sw1PerS allows us to draw similar conclusions to DAPS, but it is considerably more computationally intensive.

### 2.5 AUROC

To statistically test our hypothesis that periodicity is higher for fewer cell divisions, we use a tool from signal processing known as the *Receiver Operating Characteristic* (ROC) curve (Brown and Davis (2006)). This provides a quantitative way of comparing two distributions. Given a distribution of block periodicity scores for all cells with *n*_1_ divisions and all cells with *n*_2_ > *n*_1_ divisions, one could create a simple classifier by choosing a threshold *x* at which all blocks with periodicity scores above *x* are classified as having *n*_1_ divisions, and all blocks with scores below *x* are classified as having *n*_2_ divisions. Let *p* be the “true positive” proportion of blocks with *n*_1_ divisions which have scores above *x* and which are consequently correctly classified, and let *f* be the “false positive” proportion of blocks with *n*_2_ divisions but scores greater than *x*, which are incorrectly classified as having *n*_1_ divisions with this classifier. For *x* = 0, *f* = 0.0 and *t* = 0.0, and for *x* = 1, *f* = 1.0 and *t* = 1.0. The ROC curve is the set of all (*f, t*) pairs in between (0, 0) and (1, 1) that can be obtained by varying *x*. For the ideal case in which two distributions can be perfectly separated, there will be an *x* for which *f* = 0.0 and *t* = 1.0, but most pairs of distributions will overlap slightly. Conversely, if two distributions are exactly the same, then *f* = *t* for all *x*, which is a diagonal line. If we consider the area under the ROC curve, referred to as the “AUROC,” it is 1.0 in the former case and 0.5 in the latter case. Hence, an ROC curve further above the diagonal with an area closer to 1.0 indicates that the distributions are better separated, and an ROC curve closer to the diagonal indicates that they are more overlapping. Figure 3 shows an example of two distributions and their corresponding ROC curve. Translating our hypothesis into this framework, we expect higher AUROC values between distributions of blocks with a larger disparity between the number of divisions, and we test this hypothesis in Section 3.1.

**Fig. 3.**
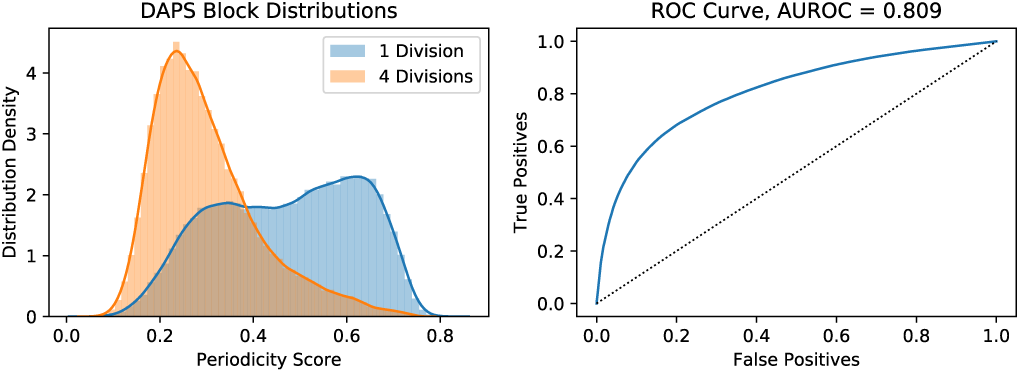
An ROC curve (right) is computed to indicate how much two distributions of periodicity scores for different numbers of divisions (left) overlap. These distributions are different, but not perfectly separated, so their ROC curve is in between the diagonal line and the upper left corner. Consequently, the area under the ROC curve (AUROC) is between 0.5 and 1.

## 3. RESULTS AND DISCUSSION

### 3.1 Localized periodicity scoring via DAPS is a good descriptor of periodicity in p53 signals

In this section, we demonstrate the capability of DAPS for summarizing the overall periodicity behavior of p53 time series. We further indicate the advantage of describing periodicity scores by a distribution, rather than by a single value (through pitch detection on the entire series).

For this purpose, we compare the pitch score distributions of two p53 time series from Reyes et al. (2018) using DAPS. At the time of the experiment, both cells were unstressed, that is, they had not been exposed to radiation. The experiment ran for a total of 5 days, with p53 measurements taken every 30 minutes. Note that in the case of cell division, the p53 time series were obtained by following one of the daughter cells (picked at random). The first cell, which did not divide during the 5 day experiment, shows mostly regular pulsing behavior in p53; see Figure 4 (top plot); the second cell, which divided 5 times during the 5 day experiment, shows mostly spontaneous pulsing behavior; see Figure 5 (top plot).

**Fig. 4.**
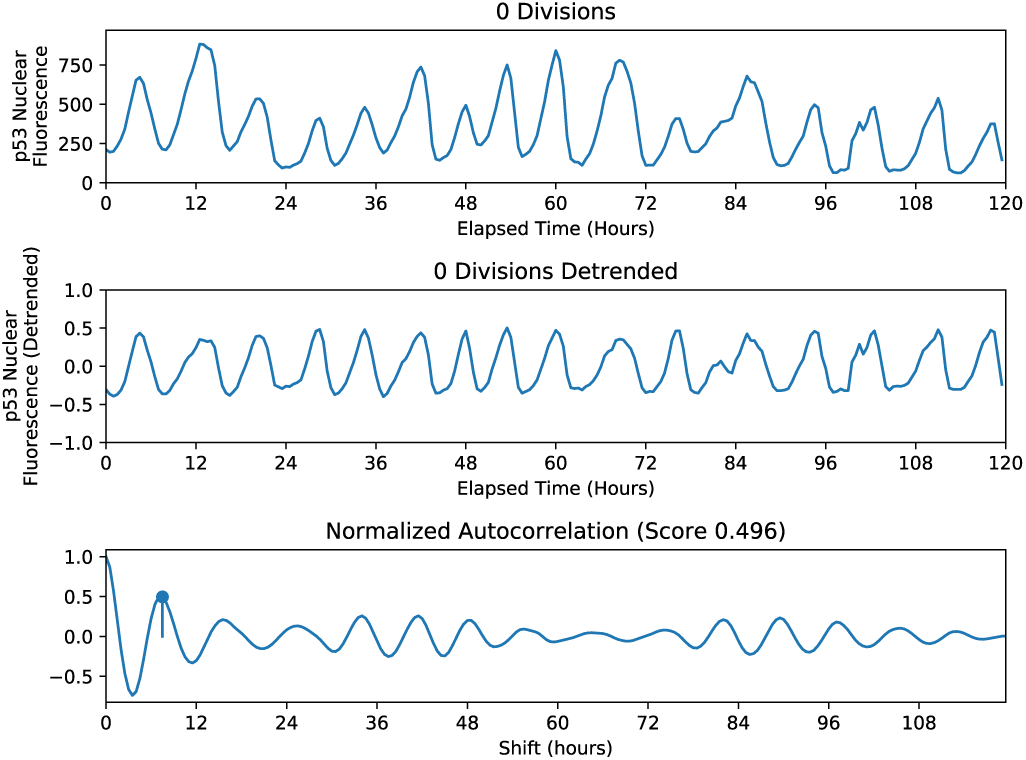
(Top plot) A p53 signal measured over 5 days (measurements every 30 minutes) of a cell that did not divide. This signal is an example of mostly regular pulsing behavior. (Center plot) The detrended version of the original signal. (Bottom plot) ACF of the detrended signal. The pitch is indicated by a blue dot and the associated pitch score is 0.496.

**Fig. 5.**
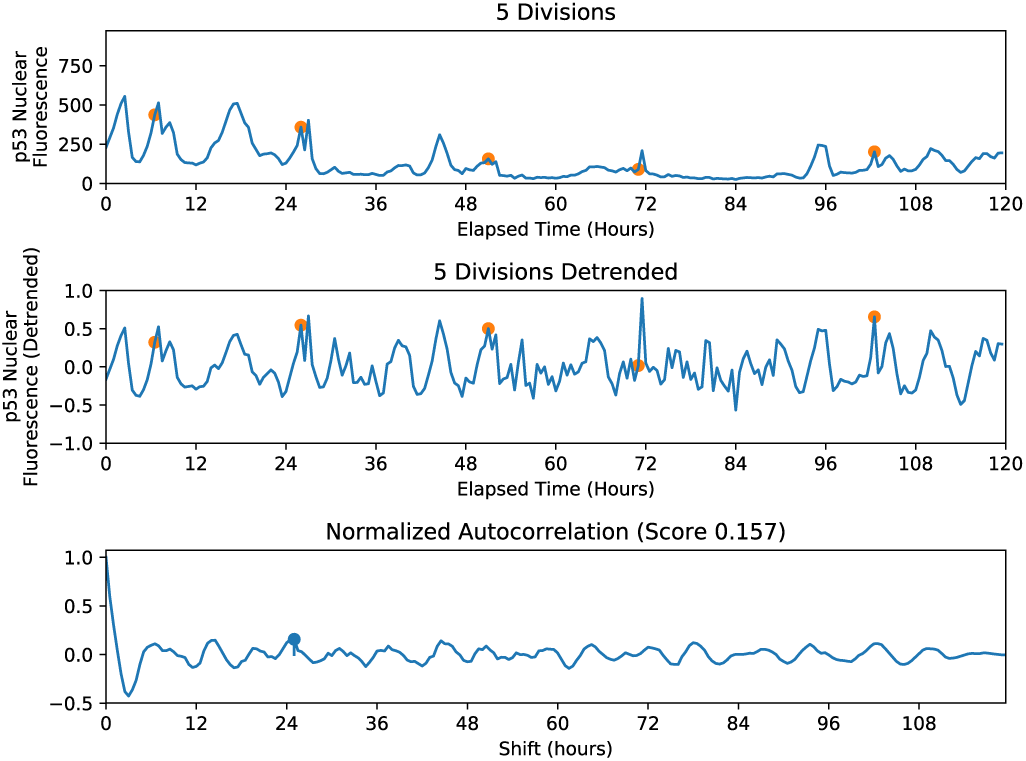
(Top plot) A p53 signal measured over 5 days (measurements every 30 minutes) of a cell which divided 5 times (division events marked with orange dots). This signal is an example of mostly spontaneous pulsing behavior. (Center plot) The detrended version of the original signal. (Bottom plot) ACF of the detrended signal. The pitch is indicated by a blue dot and the associated pitch score is 0.157.

Applying the steps of DAPS outlined in Section 2.3, we first detrend the two time series (center plots of Figure 4 and Figure 5, respectively). We then consider sliding windows of length 24 h, which is the mean duration of a cell cycle, and compute the pitch score in every 24 hour interval. This step is schematically shown in Figure 6 for the first cell (non-divider). In the final step, we summarize all periodicity scores in a distribution. Figure 7 shows the distribution of the first cell (non-divider) in blue, and of the second cell (divided 5 times) in orange. It is obvious that the blue distribution has a stronger tendency towards higher periodicity scores — a fact that is expected by comparing the two original p53 time series (top plots of Figures 4 and 5).

**Fig. 6.**
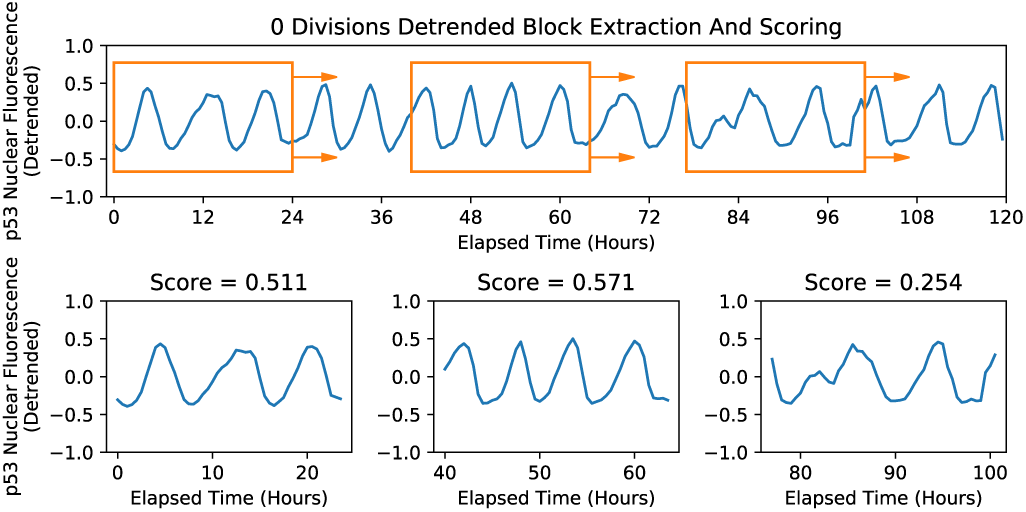
For a time series, we extract all contiguous chunks of time spanning 24 hours, which we refer to as “blocks” (shown in orange) and we provide a score individually to each block. This allows us to form a distribution of periodicity scores localized to different regions of the time series (Figure 7).

**Fig. 7.**
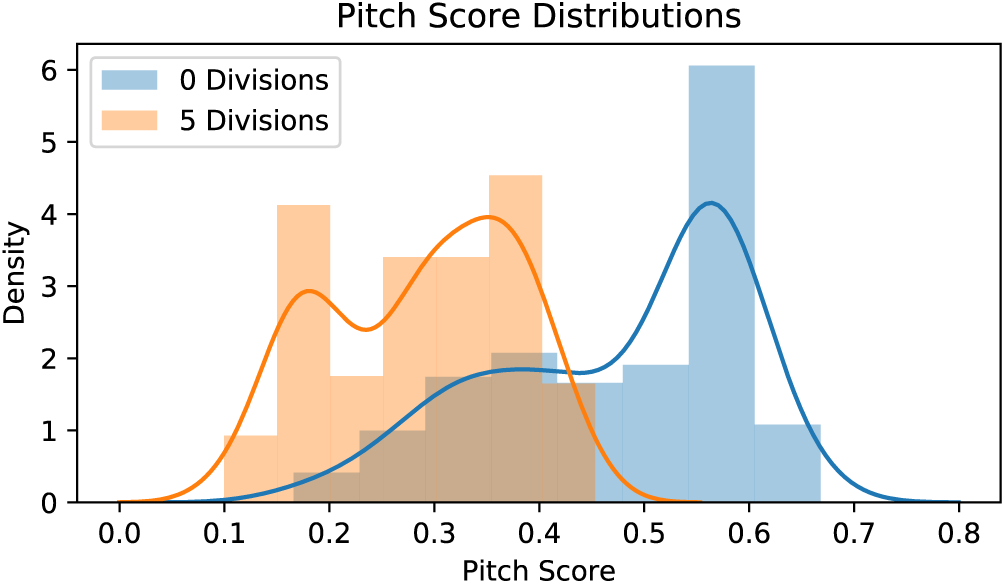
Pitch score distributions of the p53 time series of Figure 4 (blue distribution) and of the p53 time series in Figure 5 (orange distribution). The blue distribution (0 division p53 time series) has a stronger tendency towards higher periodicity scores than the orange distribution (5 division p53 time series).

We further computed the pitch score of the entire time series (the respective ACFs are shown in the bottom plots of Figures 4 and 5; the pitch is indicated by a blue dot). The first cell (non-divider) is assigned an overall pitch score of 0.496, while the second cell (divided 5 times) is assigned an overall pitch score of 0.157. While these pitch scores provide a distinction between the two signals, indicating that the first signal is indeed more periodic than the second one, the summary obtained via pitch score distributions is much more informative with respect to local periodicity changes in the time series.

### 3.2 p53 pulsing behavior under basal conditions correlates with the number of cell divisions

In this section, we establish a correlation between p53 periodicity of cells under basal conditions and number of cell division events. In particular, we show that high periodicity scoring of p53 is correlated to low number of cell divisions, and vice versa.

Through this correlation, we demonstrate that the high variability in the basal dynamics of p53 reported in the experiments of Loewer et al. (2010); Reyes et al. (2018) comes from the variability in number of cell divisions: If a cell population exhibits diverse dividing behavior, then it will also show variable pulsing in the p53 signal.

To link p53 periodicity to number cell division events, we analyze the p53 signals of unstressed cells from Reyes et al. (2018) (0 Gy data set). This data set consists of 384 cells, which had been traced over 5 days, with measurements taken every 30 minutes. In the case of cell division, the p53 time series were obtained by following one of the daughter cells (picked at random).

We apply DAPS (as outlined in Section 2.3) to all 384 p53 time series to obtain the respective pitch score distributions. To give a visual impression of how periodicity scoring is correlated to number of cell divisions, we consider six groups (one for each division type: 0 to 5 divisions), and summarize the periodicity scores of all 24 h sliding window blocks of all cells that belong to a specific division group in a distribution; see Figure 8 (top left). This figure shows that the periodicity distribution of non-dividing cells (blue distribution) has the highest tendency towards 1 (most periodic), whereas the distribution of cells which divided 4 or 5 times during the experiment (purple and brown distribution) are closest to 0 (least periodic). While there is an overlap between all of the distributions, a continuous shift from low to high periodicity is clearly visible as the number of divisions decreases.

**Fig. 8.**
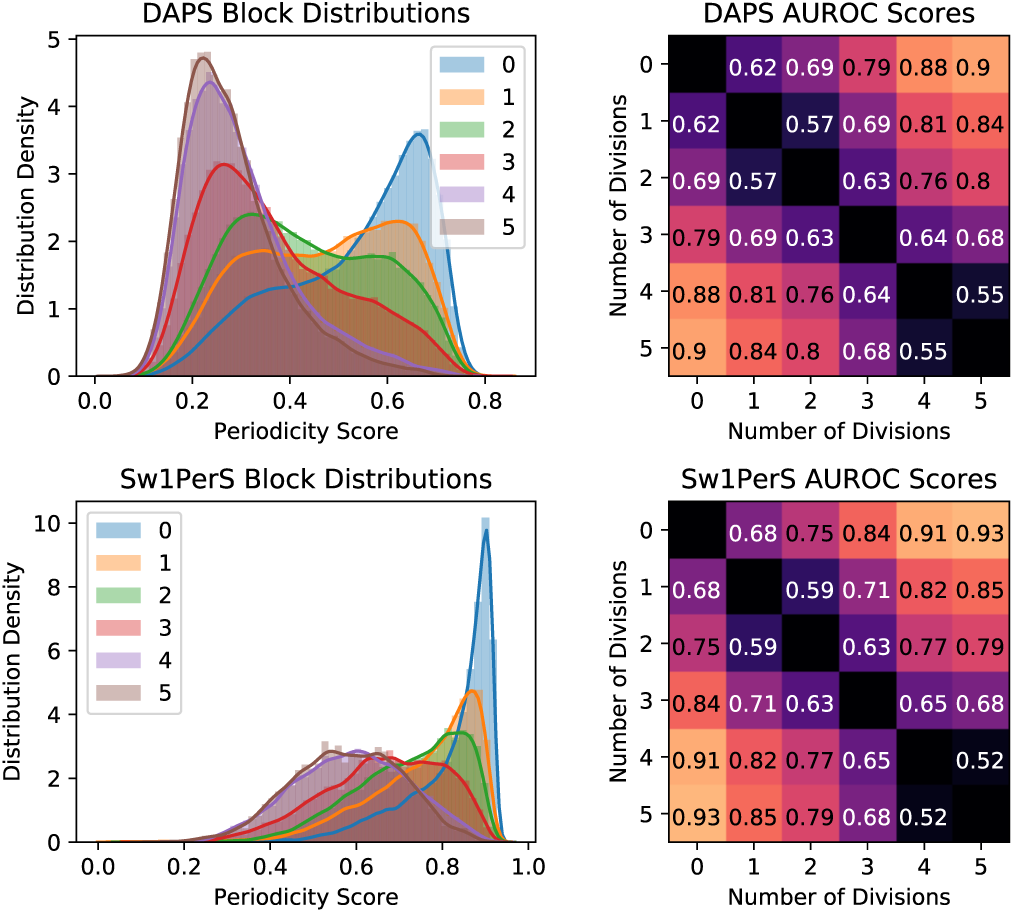
(Top and bottom left): Summary of p53 periodicity scores (computed via DAPS resp. Sw1PerS) of **all** 24 h sliding window blocks of **all cells** that belong to a specific division group in a distribution. We encounter six division groups in the data set: 0 to 5 divisions. (Top and bottom right): AUROC scores summarize how well p53 periodicity scores (computed via DAPS resp. Sw1PerS) can distinuguish between different division groups. Exact AUROC scores are superimposed in the heat map for each division pair. Heat map color code: Light color means high AUROC, dark color means low AUROC. We observe that a higher disparity between number of divisions leads to a higher AUROC score, which matches our hypothesis.

We show the capability of p53 periodicity to distinguish between the different division groups with AUROC scores (see Section 2.5). Figure 8 (top right) shows the AUROC scores between different division groups as a heat map (light color means high AUROC score, dark color means low AUROC score). We further superimpose the exact AUROC score for each division pair.

Figure 8 (top right) shows that the best distinction between two groups based on p53 periodicity is achieved when the difference of number of cell divisions is maximal: non-dividing cells can be almost perfectly separated from cells that divided 4 or 5 times (AUROC ∼ 0.9). A continuous increase of separability performance as the difference in number of cell divisions increases is also clearly visible, for example, the first column, which shows how well the group of non-dividers can be distinguished from all the other groups, goes from 0.62 (AUROC with 1-dividers) to 0.9 (AUROC with 5-dividers). A similar behavior can be observed with respect to the other division groups (other columns).

Note that all division groups can be separated by p53 periodicity, as AUROC is always greater than 0.5. Nevertheless, 4− and 5− dividers are almost identical from this point of view (AUROC= 0.55), a fact which is also visible in the distribution plot (Figure 8 top left).

We further analyzed the same data set of p53 signals with the state-of-the-art algorithm for periodicity scoring Sw1PerS (see Section 2.4). We obtained the same correlation between p53 periodicity scoring (now computed via Sw1PerS instead of DAPS) and number of cell division events; see Figure 8 (bottom row). While the AUROC scores achieved by Sw1PerS are a little better than the ones achieved by DAPS, computationally, Sw1PerS is considerably more expensive.

We note that there is a crucial difference between the data set we analyzed compared to the data set analyzed in Loewer et al. (2010). Namely, the cells in Loewer et al. (2010) are traced over 1 day to a maximum of 2 days, while the cells of Reyes et al. (2018) are traced over 5 days. The mean duration of a cell cycle is 24 hours; therefore the maximal amount of cell divisions encountered in the study of Loewer et al. (2010) is 2, while in our analysis we observe up to 5 divisions. Due to the 5 day experiment, a higher variation in number of cell divisions is achieved, which is an essential ingredient for establishing a correlation with periodicity scoring of the p53 signal.

## 4. CONCLUSION

In this paper we introduce an algorithm to score the local periodicity of time series, called Detrended Autocorrelation Periodicity Scoring (DAPS). This algorithm uses pitch detection on sliding windows of the full time series to describe the overall periodicity by local pitch scores. We apply DAPS to analyze the long-term (5 days) periodic behavior of the important tumor suppressor p53 in single cells under basal conditions. Via DAPS we are able to establish a correlation between the periodicity scoring of a p53 signal and the number of cell division events. In particular, we show that high p53 periodicity implies low number of cell divisions and vice versa. Similar results can be achieved by applying the state-of-the-art periodicity scoring algorithm Sw1PerS, which is computationally more expensive than DAPS. Through the correlation between p53 periodicity and the number of cell divisions, we are able to explain the experimentally observed high variability of p53 periodicity in unstressed cell populations by the simple observation that the number of cell division events is variable across a cell population.

In future work, we will consider a similar analysis for cells treated at various level of radiation.

## ACKNOWLEDGEMENTS

We would like to thank Professor Galit Lahav of Harvard University for making available the data used in this study.

## Footnotes

1 In the experimental p53 data we analyze in this paper, after a cell divides, one of its daughter cells is picked at random to continue with the measurement of p53.

## REFERENCES

Bello, J.P. (2011). Measuring structural similarity in music. IEEE Transactions on Audio, Speech, and Language Processing, 19(7), 2013–2025.

Box, G.E.P. and Jenkins, G.M. (1976). Time series analysis: forecasting and control. Holden-Day.

Brown, C.D. and Davis, H.T. (2006). Receiver operating characteristics curves and related decision measures: A tutorial. Chemometrics and Intelligent Laboratory Systems, 80(1), 24–38.

Chatfield, C. (2003). The analysis of time series: An introduction. Chapman & Hall/CRC texts in statistical science.

Edelsbrunner, H. and Harer, J. (2010). Computational topology: An introduction. American Mathematical Society.

Emrani, S., Gentimis, T., and Krim, H. (2014). Persistent homology of delay embeddings and its application to wheeze detection. IEEE Signal Processing Letters, 21(4), 459–463.

Frank, J., Mannor, S., and Precup, D. (2010). Activity and gait recognition with time-delay embeddings. In AAAI. Citeseer.

Geva-Zatorsky, N., Rosenfeld, N., Itzkovitz, S., Milo, R., Sigal, A., Dekel, E., Yarnitzky, T., Liron, Y., Polak, P., Lahav, G., and Alon, U. (2006). Oscillations and variability in the p53 system. Molecular Systems Biology, 2(1). doi:10.1038/msb4100068.

Herzel, H., Berry, D., Titze, I.R., and Saleh, M. (1994). Analysis of vocal disorders with methods from nonlinear dynamics. Journal of Speech, Language, and Hearing Research, 37(5), 1008–1019.

Jin, S. and Levine, A. (2001). The p53 functional circuit. Journal of Cell Science, 114(23), 4139–4140.

Khasawneh, F.A. and Munch, E. (2014). Stability determination in turning using persistent homology and time series analysis. In ASME 2014 International Mechanical Engineering Congress and Exposition. American Society of Mechanical Engineers Digital Collection.

Khasawneh, F.A. and Munch, E. (2016). Chatter detection in turning using persistent homology. Mechanical Systems and Signal Processing, 70, 527–541.

Lahav, G. (2009). Oscillations by the p53-mdm2 feedback loop. In M. Maroto and N.A.M. Monk (eds.), Cellular Oscillatory Mechanisms, 28–38. Springer New York, New York, NY. doi:10.1007/978-0-387-09794-72.

Lahav, G., Rosenfeld, N., Sigal, A., Geva-Zatorsky, N., Levine, A.J., Elowitz, M.B., and Alon, U. (2004). Dynamics of the p53-mdm2 feedback loop in individual cells. Nature Genetics, 36, 147–150.

Lev Bar-Or, R., Maya, R., Segel, L.A., Alon, U., Levine, A.J., and Oren, M. (2000). Generation of oscillations by the p53-Mdm2 feedback loop: A theoretical and experimental study. Proceedings of the National Academy of Sciences, 97(21), 11250–11255. doi: 10.1073/pnas.210171597.

Levine, A.J. (1997). p53, the cellular gatekeeper for growth and division. Cell, 88, 323–331.

Levine, A.J. (2020). P53 and the immune response: 40 years of exploration — a plan for the future. Int. J. Mol. Sci., 21(2), 541.

Loewer, A., Batchelor, E., Gaglia, G., and Lahav, G. (2010). Basal dynamics of p53 reveal transcriptionally attenuated pulses in cycling cells. Cell, 142, 89 –100.

Loewer, A., Karanam, K., Mock, C., and Lahav, G. (2013). The p53 response in single cells is linearly correlated to the number of DNA breaks without a distinct threshold. BMC Biology, 11(1), 114. doi:10.1186/1741-7007-11-114.

Ma, L., Wagner, J., Rice, J.J., Hu, W., Levine, A.J., and Stolovitzky, G.A. (2005). A plausible model for the digital response of p53 to DNA damage. Proceedings of the National Academy of Sciences, 102(40), 14266–14271. doi:10.1073/pnas.0501352102.

Martin, N. and Mailhes, C. (2010). About periodicity and signal to noise ratio - the strength of the autocorrelation function. In 7th Int. Conf. Condition Monitoring and Machinery Failure Prevention Technologies (CM & MFPT 2010). Stratford-upon-Avon, UK.

Michael, D. and Oren, M. (2003). The p53-Mdm2 module and the ubiquitin system. Seminars in Cancer Biology, 13(1), 49 –58. doi:10.1016/S1044-579X(02)00099-8.

Perea, J.A., Deckard, A., Haase, S.B., and Harer, J. (2015). SW1Pers: Sliding windows and 1-persistence scoring; discovering periodicity in gene expression time series data. BMC Bioinformatics, 16(1), 257. doi: 10.1186/s12859-015-0645-6.

Perea, J.A. and Harer, J. (2015). Sliding windows and persistence: An application of topological methods to signal analysis. Foundations of Computational Mathematics, 15(3), 799–838. doi:10.1007/s10208-014-9206-z.

Reyes, J., Chen, J.Y., Stewart-Ornstein, J., Karhohs, K.W., Mock, C.S., and Lahav, G. (2018). Fluctuations in p53 signaling allow escape from cell-cycle arrest. Molecular Cell, 71, 1–11.

Stam, C.J. (2005). Nonlinear dynamical analysis of eeg and meg: review of an emerging field. Clinical Neurophysiology, 116(10), 2266–2301.

Stewart-Ornstein, J. and Lahav, G. (2017). p53 dynamics in response to DNA damage vary across cell lines and are shaped by efficiency of DNA repair and activity of the kinase ATM. Science Signaling, 10(476). doi: 10.1126/scisignal.aah6671.

Takens, F. (1981). Detecting strange attractors in turbulence. In Dynamical systems and turbulence, Warwick 1980, 366–381. Springer.

Venkataraman, V. and Turaga, P. (2016). Shape descriptions of nonlinear dynamical systems for video-based inference. IEEE transactions on pattern analysis and machine intelligence.

Vlachos, M., Yu, P., and Castelli, V. (2005). On periodicity detection and structural periodic similarity. In Proceedings of the Fifth SIAM International Conference on Data Mining. Newport Beach, CA.

Vogelstein, B., Lane, D., and Levine, A.J. (2000). Surfing the p53 network. Nature, 408, 307–310.

